# CEMIG: Prediction of the *cis*-regulatory motif using the De Bruijn graph from ATAC-seq

**DOI:** 10.1101/2023.05.26.542440

**Authors:** Yizhong Wang, Yang Li, Cankun Wang, Qin Ma, Bingqiang Liu

## Abstract

Sequence motif discovery algorithms identify novel DNA patterns with significant biological roles, such as transcription factor (TF) binding site motifs. Chromatin accessibility data, accumulated through assay for transposase-accessible chromatin with sequencing (ATAC-seq), has enriched resources for motif discovery. However, computational efforts in ATAC-seq data analysis mainly target TF binding activity footprinting rather than motif prediction. Here, we introduce CEMIG, an algorithm predicting and characterizing TF binding sites, leveraging the De Bruijn and Hamming distance graph models. Evaluation of 129 ATAC-seq datasets from the Cistrome Data Browser suggests that CEMIG outperforms three widely used methods using four metrics. It is noteworthy that CEMIG is employed to predict cell-type-specific and shared TF motifs in GM12878 and K562 cells, facilitating comprehensive gene expression and functional genomics analysis.

## Introduction

Regulatory proteins such as transcription factors (TFs) and RNA binding proteins modulate both transcriptional and post-transcriptional regulation by binding to DNA and RNA^1-9^. Computational tools have utilized Chromatin immunoprecipitation and high-throughput sequencing (ChIP-seq) to *de novo* predict protein binding sites, or motifs, yielding a plethora of genome-wide protein-DNA and protein-RNA interactions. However, ChIP-seq, while identifying DNA fragments linked to specific TFs or histone marks, yields broader and less precise peaks. Assay for transposase-accessible chromatin with sequencing (ATAC-seq), a prevalent protocol for the genome-wide identification of open chromatin, leverages cleavage enzymes (Tn5) that recognize and cleave open chromatin regions^10^. Sequencing and alignment of these fragments facilitate open chromatin detection by identifying genomic intervals with abundant reads. ATAC-seq, not requiring prior knowledge of TF binding and capable of handling rare cell subpopulations with minimal background noise, emerges as a valuable complementary method for studying gene regulation^10^.

Over the past decade, ATAC-seq-based TF binding analysis has chiefly centered on footprinting, i.e., identifying ATAC regions experiencing a drop in cleavage events due to TF binding^11,12^. A common approach, ATAC-seq footprinting, identifies open chromatin regions and predicts TF binding sites (TFBSs). It rests on the premise that TF binding to a particular DNA sequence induces chromatin structure alterations detectable by ATAC-seq. However, ATAC-seq footprinting falls short in directly pinpointing specific TFBSs. The technique suggests TF binding site presence by scanning for characteristic ATAC-seq read density decreases, known as ‘footprints’, in open chromatin regions. While ATAC-seq can locate TF-occupied open chromatin regions and offer some TF binding site location insights, it fails to differentiate between varying TF binding events. This limitation arises from the footprint specificity issue; the footprint of a TF may resemble those of other TFs or DNA-binding proteins. Consequently, ATAC-seq footprinting struggles to precisely identify specific TFBSs and discern the binding specificity of a TF for different DNA sequences. Thus, *de novo* motif finding based on ATAC-seq peaks remains necessary.

Several existing computational methods, such as BioProspector, XXmotif, MEME-ChIP, and STREME, offer solutions. BioProspector employs Gibbs sampling, a probabilistic algorithm, to pinpoint motifs statistically overrepresented in input sequences, without initial assumptions about motif size distribution or pre-defined background models^13^. XXmotif, a position-weighted matrix (PWM) based method, optimizes PWMs by directly minimizing the *P*-values of enrichment, ensuring a reliable and sensitive tool for gene regulatory network analysis^14^. MEME-ChIP^15^, a tool for ChIP-seq data analysis to identify TF binding motifs, employs two motif-finding algorithms, MEME and DREME (replaced by STREME), to identify overrepresented sequence motifs^7,16^. The MEME algorithm depicts input sequences as a blend of Markov models, facilitating the identification of both exact and degenerate motifs^15^. This algorithm boasts extensive customization, letting users determine parameters like the maximum motif width, the count of motifs to be detected, and the minimum motif incidence in the input sequences. Meanwhile, STREME, a motif-finding algorithm, recognizes overrepresented sequence motifs in DNA or RNA sequences^16^. It discerns motifs by building a suffix tree from the input sequences, enumerating all substrings appearing at least *k* times in the input, evaluating motif candidates for statistical relevance and biological significance, and linking the identified motifs to known TF binding site databases.

Despite their robustness, these methods may not be suitable for ATAC-seq due to the inherent differences from ChIP-seq. MEME assumes that input sequences possess shared, yet unidentified, TF motifs. These techniques assess TF motifs via statistical tests on *k*-mer frequencies between input and background sequences. However, ATAC-seq data might have a higher prevalence of non-specific background noise compared to ChIP-seq or promoter sequences, presenting challenges in the accurate identification of true motifs^16^. Furthermore, ATAC-seq data may exhibit a lower signal-to-noise ratio than ChIP-seq or promoter sequences, complicating the accurate determination of enriched regions and the differentiation between true and false positive motifs. STREME, in particular, operates on the presumption that motifs reside in specific regions of the input sequences, such as promoter or enhancer zones. Yet, ATAC-seq data fails to offer precise promoter or enhancer locations in the genome, thereby restricting the accuracy of motif identification via STREME^16^. Thus, there is a clear demand for novel computational approaches designed specifically for motif detection in ATAC-seq data.

In this article, we introduce CEMIG (<u>C</u>is r<u>E</u>gulatory <u>M</u>otif Influence using De Bruijn <u>G</u>raph), an algorithm based on the De Bruijn Graph (DBG), to predict and characterize TF motifs^17^. CEMIG utilizes graph clustering and path detection on DBG to achieve this task. Specifically, it assesses and ranks all *k*-mers from ATAC-seq data using *P*-values derived from a Poisson distribution. Subsequently, we construct two graph models: the Hamming distance graph and DBG, based on these *k*-mers. The Hamming distance graph illustrates the Hamming distances between *k*-mers. In DBG, vertices symbolize *k*-mers, with directed edges linking two *k*-mers if the (*k* − 1) suffix of one *k*-mer matches the (*k* − 1) prefix of the other. *k*-mers that exhibit proximity, as measured by Hamming distance, are grouped on the Hamming distance graph. In an updated DBG, vertices representing *k*-mers within the same group consolidate into a single cluster vertex. This updated DBG is then deployed to discover paths through extension, beginning from the cluster vertex. Each path found is segmented into one or more motifs according to the overlap of occurrences upstream or downstream relative to the occurrences of the starting cluster vertex.

Distinguished from MEME, CEMIG does not operate under the presumption that a significant portion of the input sequences contains identical, unknown TF motifs. Moreover, the effectiveness of CEMIG remains intact even in the presence of a low signal-to-noise ratio, owing to its dual strategy of strengthening true signals. This strategy consists of clustering alike *k*-mers and building motifs based on DBG paths rather than merely using *k*-mers. Even if certain false positive *k*-mers achieve significant *P*-values, the likelihood of their inclusion in a DBG path remains minimal. Additionally, CEMIG offers the convenience of autonomously determining motif lengths, eliminating the need for users to provide an interval of candidate motif lengths. As a result, CEMIG circumvents the pitfalls of low signal-to-noise ratios and specific parameter influences, thus enhancing its overall robustness.

In this article, we first describe the CEMIG algorithm with algorithm design framework and rationale (see technical details in **Methods**) and present experimental results comparing its performance with BioProspector, MEME-ChIP, and XXmotif on ATAC-seq data in Cistrome Data Browser composed of different numbers of peaks^18^. Experimental results indicate that our proposed CEMIG can accurately predict sequences that contain TFBSs and outperform the other three methods evaluated by precision, specificity, accuracy (ACC), and the area under the precision-recall curve (PRC)^2^. Beyond these observations, we also explore the ability of CEMIG to predict cell-type-specific and shared TF binding activities in GM12878 and K562, showing that four important TFs possess different binding specificities in GM12878 and K562. TF binding specificities also underlie the differential expression of genes closest to TF binding peaks in the two cell types, and thus divergence in functionalities depicted by enriched gene ontology (GO) terms and KEGG pathways^19,20^.

## Results

### Framework of CEMIG

The fundamental framework of CEMIG can be clarified as follows (see **Fig. 1**): Initially, CEMIG ingests the input sequences in either BED or FASTA formats while concurrently estimating the likelihood of nucleotide occurrences based on a zero, first, and second-order Markov model. This information aids in determining the expected frequencies of *k*-mer (*k* = 6) within the imported sequences (**Fig. 1a**). By employing a Poisson distribution model to represent *k*-mer frequencies, CEMIG can compute the associated *P*-values^21,22^. These *P*-values, in turn, are utilized to filter and rank notable *k*-mers, which form the building blocks of potential motifs. In the subsequent step, CEMIG arranges all *k*-mers in a descending sequence, based on their negative logarithm *P*-values, and segments them into three subsets: top 100 (most significant), the first half (moderately significant), and the second half (insignificant). This distribution aids in the construction of both the Hamming distance graph and DBG. For the Hamming distance graph, the algorithm leverages the first half of *k*-mers. In this structure, vertices symbolize *k*-mers, edges correspond to *k*-mer pairs with a Hamming distance not exceeding two, and the weight of each edge mirrors its Hamming distance (**Fig. 1b**). The DBG, on the other hand, is created using all *k*-mers as vertices. It establishes a directed edge connecting one *k*-mer to another if the (*k* − 1) prefix of the first *k*-mer coincides with the (*k* − 1) suffix of the second. The weight assigned to each directed edge mirrors the frequency of the (*k* + 1) -mer, which results from overlapping the initial (*k* − 1) nucleotides of the first *k*-mer with the concluding (*k* − 1) nucleotides of the second *k*-mer (**Fig. 1b**). In the third phase, CEMIG conducts clustering on the Hamming distance graph, followed by the construction of a second directed graph (digraph) by merging the vertices within each cluster into a singular vertex on the DBG. This clustering process involves the identification of maximal Independent Sets (IS) based on the subset of the top 100 *k*-mers and a greedy algorithm that maximizes an objective function on the Hamming distance graph (**Fig. 1c**). Following the DBG, CEMIG assembles the second digraph by substituting the *k*-mer-denoting vertices within the same cluster with the corresponding cluster (cluster vertex). It then updates edge weights by accumulating the weights of all edges connecting *k*-mers within a specific cluster to their respective neighbors. Fourth, CEMIG ranks all clusters in decreasing order of the objective function values and greedily extends a path in two directions from the top-ranked cluster vertex until reaching the termination condition (**Fig. 1d**). Each path acts as a seed to construct motifs based on: (*i*) the sub-path upstream of the starting cluster vertex, and (*ii*) the sub-path downstream of the starting cluster vertex.

**Fig. 1.**
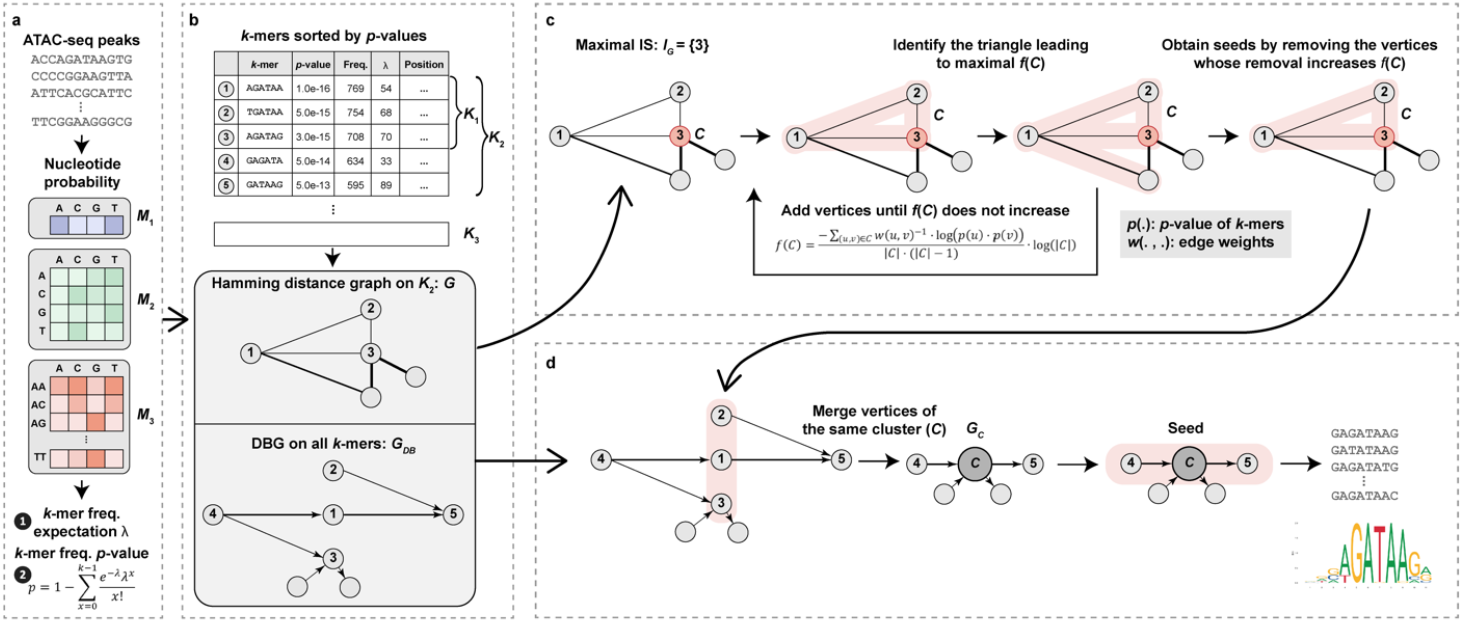
Visual representation of the CEMIG framework. **a** Calculation of *k*-mers frequencies in the background using Markov models. **b** Building Hamming distance (*G*) and De Bruijn (*G*_*DB*_) graphs with *k*-mers. **c** Grouping *k*-mers via clustering on *G*, resulting in the graph *G*_*C*_ which consolidates *k*-mers of the same cluster on *G*_*DB*_. **d** Discovery of motifs through path extensions on *G*_*C*_

### Comparison of motif discovery algorithms on ATAC-seq data

The efficacy of a motif discovery algorithm in processing an ATAC-seq dataset hinges on its ability to discern sequences harboring at least one TFBS. This challenge can be viewed as a sequence classification task, evaluated via precision, specificity, ACC, and PRC. We sourced 129 human ATAC-seq datasets from the Cistrome Data Browser (**Supplementary Table 1**)^18^. In every dataset, the sequences were derived from ATAC-seq footprints, all of which were positive sequences tagged with ‘1’^11^. By randomly rearranging all bases within each positive sequence using the MEME suite, we generated a corresponding negative sequence tagged with ‘0’ (**Fig. 2a**)^15,23^. Upon amalgamation of positive and negative sequences for each dataset, we compiled benchmarking data comprising between 848 and 692,368 sequences. These datasets were then stratified into 15 ascending groups based on sequence count (**Figs. 2b-2p**).

**Fig. 2.**
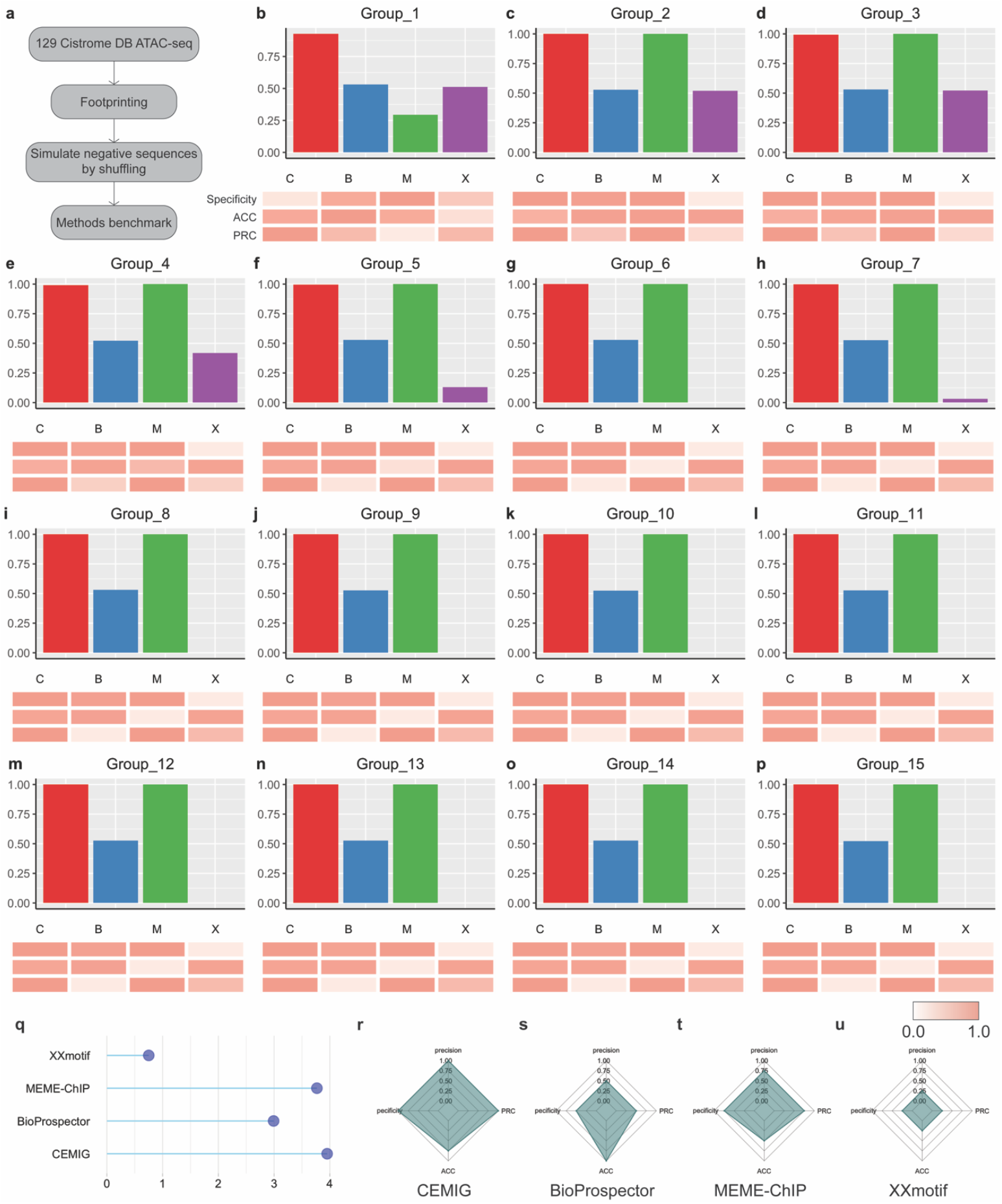
Performance evaluation of CEMIG and alternative motif discovery methods using the Cistrome Data Browser. **a** Illustration of benchmark data and evaluation strategy. **b-p** Bar charts: Display of precision comparisons across 15 grouped datasets. C, B, M, and X symbolize CEMIG, BioProspector, MEME-ChIP, and XXmotif, respectively. Heatmaps: Comparative display of four algorithmic performances in specificity, ACC, and PRC, respectively (normalized over the four algorithms). **q** Lollipop chart: Aggregate scores of each algorithm across the 15 groups of 129 datasets. **r-u** Spider charts: Average performance scores of each algorithm in precision, specificity, ACC, and PRC.

The bar plots reveal that CEMIG consistently attains the highest precision values across all datasets, although MEME-ChIP presents a comparable precision performance. This similarity may stem from the quality control steps employed in the MEME algorithm within MEME-ChIP, such as prioritizing the central 100 bp regions of each sequence^24^. Despite its stable precision over a range of dataset sizes, BioProspector shows enhanced precision in datasets with a relatively smaller number of sequences (*n* ≤ 231,120), outpacing XXmotif, which exhibits better performance with larger datasets (**Figs. 2b-2f**). However, XXmotif often fails to produce results after a 24-hour run on larger datasets (**Figs. 2g-2p**). Regarding specificity, or the ability to discern false negatives, CEMIG, BioProspector, and MEME-ChIP offer similar results, while XXmotif tends to produce more false negatives (**Figs. 2g-2p**). When evaluating ACC, a comprehensive measure of correct predictions versus total predictions, XXmotif outperforms MEME-ChIP. This reversal might occur because XXmotif is more prone to detecting a few false negatives. The edge gained by XXmotif could be attributed to its *P*-value optimization process during discriminative motif discovery^14^. Comparatively, in binary classification assessment, reflected in PRC, the performance of BioProspector dwindles. Notably, this reduction becomes more pronounced as the number of sequences escalates. Although BioProspector remains efficient in analyzing small datasets and even maintains its speed on larger ones, the incidence of both false positive and false negative results tends to increase alongside data size^1^. CEMIG and MEME-ChIP, meanwhile, exhibit roughly comparable performance when assessed by PRC.

To present the overall efficiency of an algorithm more intuitively, we normalize and aggregate the four individual scores, resulting in a single composite score (**Fig. 2q**). CEMIG and MEME-ChIP demonstrate better overall performance than BioProspector and XXmotif. In addition, we compare the performance of the four algorithms based on the ranks of their scores. Specifically, a score of 1.00 is assigned to the algorithm that possesses the best performance, and a score of 0.25 is given to the method that has the worst performance. We calculated the average scores for each algorithm across the 15 groups of the 129 datasets to gain insight into their overall performance and respective strengths and weaknesses (**Figs. 2r-2u**). CEMIG has the highest average score regarding each of the four scores, whereas MEME-ChIP achieves high precision, specificity, and PRC. BioProspector possesses outstanding ACC among the four metrics. XXmotif, however, does not deliver ideal results, which may result from the fact that it did not output results within a limited period (24 hours). To affirm the proficiency of CEMIG in discerning motifs from ATAC-seq data, we employed TOMTOM for a comparative analysis between motifs identified by CEMIG and those curated in HOCOMOCO v.11 (**Supplementary Data 1**). Moreover, we also scrutinized the effectiveness of CEMIG in uncovering motifs from 28 ChIP-seq datasets through comparison with MEME-ChIP and DESSO (**Supplementary Table 2** and **Supplementary Fig. 1**).

### Prediction of cell-type-specific and shared TF binding

We appraised the predictive performance of CEMIG in discerning shared and cell-type-specific TF binding from an ATAC-seq dataset of GM12878 and K562 cells^25^. These cells, known for their extensive reference epigenomes, serve as invaluable resources for researchers delving into epigenetic regulation. In our study, CEMIG enables the examination of various TFs binding shared and cell-type-specific peaks in the two cell lines, their differing roles in gene regulation, and functionalities signified by pathways. Of the 1,700,884 ATAC-seq peaks, we identified 230 GM12878-specific, 32,457 K562-specific, and 579 shared peaks bound by 55 distinct TF motifs (**Figs. 3a-3d, 3i-3k**). We found that 25 of these motifs significantly matched (*Q*-values < 0.05) to 21 reference TF motifs of humans in HOCOMOCO v.11, as evaluated by TOMTOM (**Supplementary Table 1** and **Supplementary Fig. 2**). We applied the same cutoffs for *P*-value and log-fold changes as the previous study to filter cell-type-specific and shared peaks^25^. Although there are more K562-specific peaks bound by various TFs compared to GM12878, GM12878-specific peaks bound by TFs exhibited wider ranges in both *P*-value and log-fold change (**Figs. 3a-3d, 3i-3k**). We sought to elucidate the impact of differential accessibility to gene expression in the two cell types by identifying genes closest to each peak and extracting its gene expression values, quantified by fragments per kilobase of exon per million mapped fragments (FPKMs) (**Figs. 3e-3h, 3l-3n**). Given that the K562 cell line generally exhibits lower gene expression levels than GM12878 across a wide range of genes^26^, we omitted the comparison using RNA-seq data in K562 to ensure clear contrasts. As indicated by the boxplots, genes for cell-type-specific peaks bound by the four representative TFs are differentially expressed, yielding *P*-values < 0.05.

**Fig. 3.**
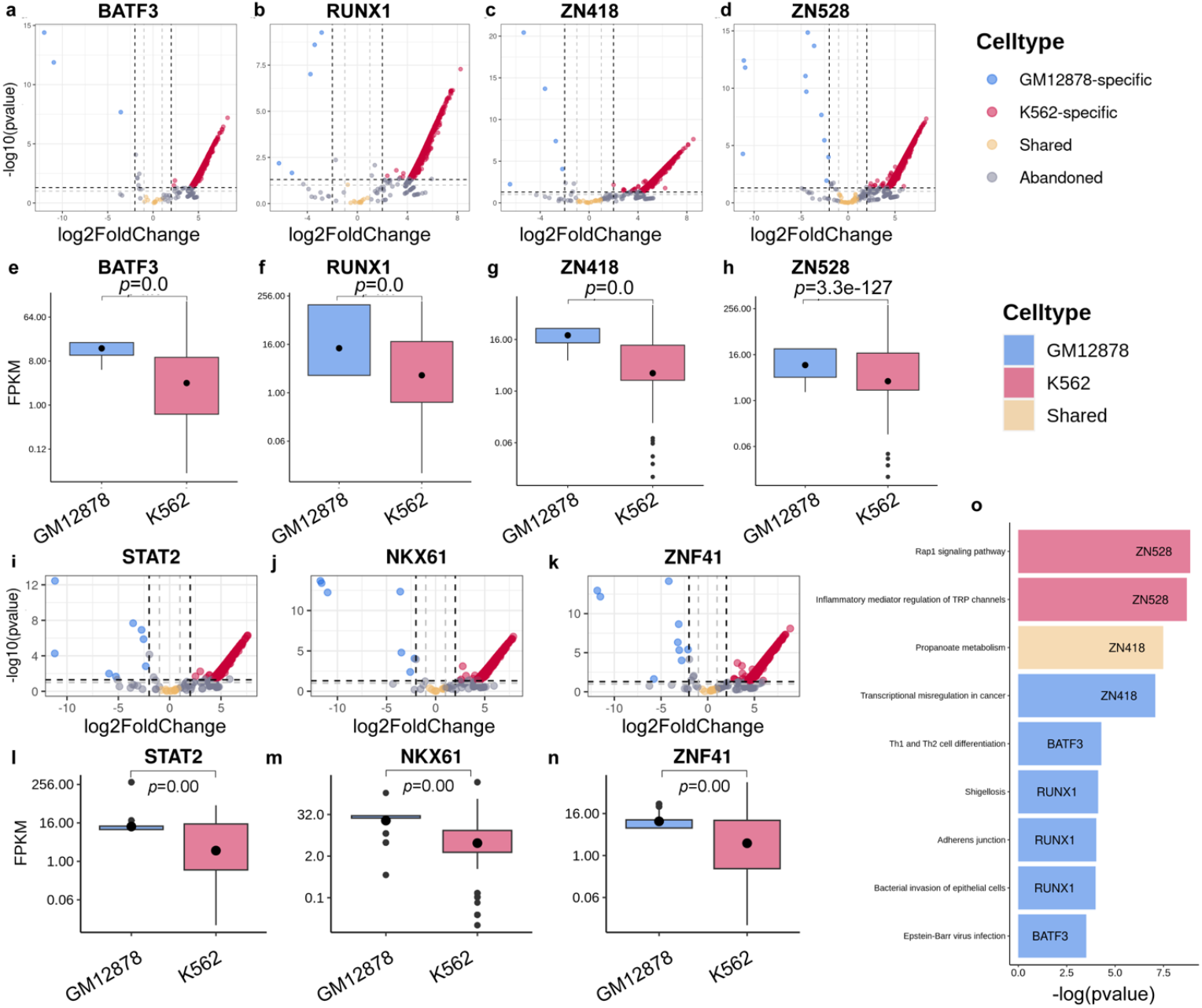
Examination of Cell-Type-Specific and Shared TF Binding. Panels **a-d, i-k** display the binding peak distribution for BATF3, RUNX1, ZN148, ZN528, STAT2, NKX61, and ZNF41. Peaks with a *Q*-value < 0.05 and a log2 fold-change less than 2 are designated as GM12878-specific binding sites. Those with a *Q*-value > 0.05 and a log2 fold-change > 2 are identified as K562-specific binding sites, and those with a *Q*-value > 0.1 and the absolute value of the log2 fold-change < 1 are recognized as shared binding sites. Red labeling denotes discarded peaks. Panels **e-h, l-n** illustrate cell-type-specific gene expression (RNA-seq) of genes proximal to binding peaks of BATF3, RUNX1, ZN148, ZN528, STAT2, NKX61, and ZNF41 between GM12878 and K562. Wilcoxon rank sum test *P*-values are indicated above the boxes. Panel **o** visualizes pathway enrichment of target genes nearest to GM12878-specific, K562-specific, and shared peaks bound by BATF3, RUNX1, ZN418, and ZN528.

To comprehend the biological functionalities of these gene sets, we undertook functional genomics analyses, evaluating their enrichment against GO terms and KEGG pathways (**Fig. 3o** and **Supplementary Table 3**). While GM12878 is not a cancer cell line, it is frequently deployed as a model system for investigations into gene regulation, intricate signaling networks, and the epigenetic groundwork behind cancer progression. Cancer misregulation is typically typified by alterations in cellular signaling pathways and gene expression programs^27^, potentially leading to rampant cell growth and proliferation, evasion of cell death, and the ability to invade and metastasize to other tissues^28^. We examined various TFs, such as BATF3, a member of the AP-1 family primarily found in dendritic cells^29^. Induction of BATF3, activated by the canonical NF-κB pathway, regulates Epstein-Barr virus (EBV) gene expression^30,31^. This induction, in response to EBV infection of GM12878 cells, indicates the potential contribution of BATF3 to the development of EBV-related diseases^32^. Although its presence is unusual in B-lymphoblastoid cells like GM12878, the low-level expression of BATF3 hints at its possible involvement in immune regulation^32^. The GM12878 B cells participate significantly in adaptive immunity by producing antibodies and presenting antigens to T cells. We also analyzed RUNX1, a TF crucial for hematopoiesis regulation. Expressed mainly in hematopoietic cells, it regulates innate immune responses^33^, including antimicrobial peptide production, neutrophil activation, and macrophage suppression^34,35^. Additionally, RUNX1 may regulate bacterial invasion in GM12878 epithelial cells^36^, with potential implications in diseases like Shigellosis^37^. This TF also aids cell adhesion and migration regulation in GM12878 cells^38,39^. Genes proximal to GM12878-specific, K562-specific, and shared peaks bound by ZN418 were enriched in cancer-related transcriptional misregulation and propanoate metabolism pathways^40^. ZN418, listed among the top 100 most expressed TFs in GM12878 by the ENCODE project^41^, is speculated to participate in cancer progression due to its downregulation^40^. While there is no direct evidence connecting ZNF418 to propanoate metabolism in GM12878 and K562, Zinc Finger (ZNF) proteins are known to regulate genes involved in lipid metabolism and hypoxia response in these cells^42,43^. Lastly, ZN528, or ZNF protein 528, is another TF implicated in regulating various cellular processes like gene expression, cell proliferation, and differentiation^44^. Its role is essential in the activation of Rap1 signaling in K562 cells, affecting the proliferation and survival of these cells^45,46^. Additionally, ZN528 contributes to the expression regulation of the interleukin-1 receptor (IL-1R), a cytokine receptor linked to the regulation of inflammatory mediators^47^.

## Discussion

Investigation into the computational prediction of TFBSs from ATAC-seq has not received as much attention as predictions derived from ChIP-seq. The latter identifies DNA fragments bound by specific TFs, but requires pre-existing knowledge of these target TFs. Despite the existence of several computational methods for TF binding analysis on ATAC-seq, and their notable performance, they predominantly focus on footprinting analysis rather than the de novo prediction of TF motifs and TFBSs. In this manuscript, we introduce a novel algorithm, CEMIG, built on the DBG model for TF motif prediction. A comprehensive evaluation of CEMIG is carried out, benchmarked against four widely-used motif-finding tools: BioProspector, MEME-ChIP, and XXmotif. Further, we contrast the TF motifs detected with reference motifs in the HOCOMOCO v.11 database, using the TOMTOM tool. PWMs of motifs identified by CEMIG demonstrate a substantial similarity (judged by *Q*-value) with the curated reference motifs. To further showcase the prowess of CEMIG, it is deployed to predict TFBSs on GM12878-specific peaks, K562-specific peaks, and peaks shared by both cell types. As anticipated, the cell-type-specific TFBSs indicate differential gene expression adjacent to these sites. Further, functional genomics analysis performed on these gene sets uncovers distinct GO terms and pathways across different cell types or binding TFs, highlighting the versatility and utility of CEMIG in genomic and epigenomic research.

## Methods

### Data acquisition

Datasets from ATAC-seq (*n* = 129) and ChIP-seq (*n* = 28) were compiled for benchmarking, gathered from the Cistrome Data Browser and ENCODE databases^18,41^. Peak data (in BED format) for each dataset were extracted from these sources, with alignment to either hg38 or hg19 human reference genomes accordingly. For the focused exploration of cell-type-specific and shared TF binding, binary alignment map (BAM) files for ATAC-seq in GM12878 and K562 cell lines were retrieved from ENCODE^41^. Each cell line was represented by two replicate files. Additionally, FPKM matrices for RNA-seq in the two cell lines were procured from ENCODE. Comprehensive details on all the ATAC-seq and ChIP-seq datasets utilized in this study can be found in **Supplementary Table 1** and **Supplementary Table 2**.

### Estimation of k-mer frequencies in the background employing Markov models

CEMIG accepts sequences in either BED or FASTA format and consolidates them into a set, *S* = {*S*_1_ …, *S*_*n*_}, where each sequence has corresponding lengths *l*_1_ *l*_2_, …, *l*_*n*_ (**Fig. 1a**). In instances where the BED format is the input, CEMIG concurrently imports the reference genome file (in FASTA format), transforming the ATAC-seq peak intervals from the BED file into FASTA sequences. Subsequently, it constructs matrices *M*_1_, *M*_2_, and *M*_3_, of dimensions 1 × 4, 4 × 4, and 16 × 4, respectively, grounded in zero, first, and second-order Markov models. To clarify, *M*_1_ represents the probabilities of individual nucleotide occurrences in the input sequences, *M*_2_ anticipates the probabilities of nucleotide occurrences considering the preceding nucleotide, and *M*_3_ signifies the probabilities of nucleotide appearances accounting for the preceding dinucleotides. Utilizing a Poisson distribution, CEMIG estimates the expected frequency of a *k*-mer (*t* = *a*_1_*a*_2_, …, *a*_*k*_), where *k* equals 6, in the given input sequences^21,22^.

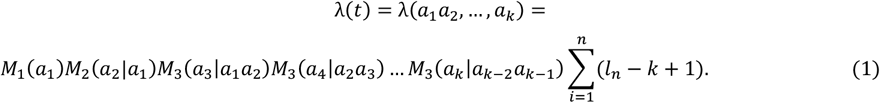

Utilizing the frequency of k-mer *t* (*x*) as input, CEMIG computes the *P*-value for the *k* –mer *a*_1_*a*_2_, …, *a*_*k*_ as

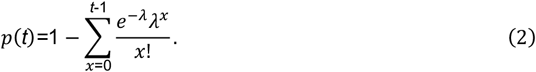

### Construction of graph models

#### Classification of k-mers

CEMIG records the occurrences of each *k*-mer in the input sequences and arranges them in descending order based on the negative logarithm *P*-values. Subsequently, CEMIG identifies the top 100 *k*-mers as the most significant set (*K*_1_), while the first half of all *k*-mers constitutes the moderately significant set (*K*_2_). The remaining half beyond *K*_2_is referred to as *K*_3_, representing the collection of insignificant *k*-mers (**Fig. 1b**).

#### Hamming distance graph construction

CEMIG generates a Hamming distance graph, denoted as *G*, where the vertices correspond to *k*-mers within the set *K*_2_(**Fig. 1b**). When the Hamming distance between two *k*-mers is less than or equal to two, CEMIG establishes an edge between the respective vertices. The weight assigned to each edge is equal to the Hamming distance.

#### DBG creation

CEMIG proceeds to construct a DBG, denoted as *G*_*DB*_ by considering all *k*-mers from the union of sets *K*_2_ and *K*_3_ as vertices (**Fig. 1b**). To establish directed edges in the graph, CEMIG identifies pairs of *k*-mers that exhibit a (*k* − 1)-nucleotide overlap between their head and tail regions. For instance, if the *k*-mers A<u>GCTAG</u> and <u>GCTAG</u>C share an overlap of size *k* − 1, a directed edge is added from A<u>GCTAG</u> to <u>GCTAG</u>C, with the edge weight corresponding to the frequency of the (*k* + 1)-mer, A<u>GCTAG</u>C, within the input sequences. It is worth noting that CEMIG applies this procedure to both the original sequences and their reverse complements, aligning with the convention followed by most motif-finding methodologies.

### Graph clustering on Hamming distance graph

CEMIG executes a clustering process on the Hamming distance graph *G*, involving two distinct steps: identification of the maximum independent set (IS) and construction of *k*-mer clusters (**Fig. 1c**).

#### Maximum IS identification

To identify the maximum independent set (IS), CEMIG employs a greedy algorithm that proceeds as follows approximately:

1. Arrange the *k*-mers in the set *K*_2_ in descending order based on their negative logarithm *P*-values,
2. Initialize the maximum IS, denoted as *I*_*G*_, as an empty set,
3. Examine each vertex, *ν*, in the set *K*_1_\*I*_*G*._ individually,
4. If vertex *ν* is not adjacent to any vertex in *I*_*G*_, include it in *I*_*G*_,
5. If vertex *ν* is adjacent to at least one vertex in *I*_*G*_, skip it and proceed to the next vertex,
6. Once all vertices have been evaluated, return the set *I*_*G*_ as the maximum IS.

CEMIG operates through an iterative process of adding vertices to the IS in a manner that preserves the independence of *I*_*G*_. To maintain independence, CEMIG examines the adjacency of each vertex to the current IS, thereby preventing the inclusion of vertices that would compromise its independence. Although CEMIG does not guarantee the identification of the true maximum IS, it frequently yields a satisfactory approximation solution.

#### k -mers cluster construction

CEMIG identifies *k*-mer clusters based on each vertex in *I*_*G*_ as follows.

1. Sort the vertices in *I*_*G*_ in descending order according to negative logarithm *P*-values,
2. Initialize an empty set as the cluster set,
3. For each vertex *ν* in *I*_*G*_
  a. Initialize the current cluster *C* = {*ν*},
  b. Find all neighbors of *ν*, denoted by *N*_*G*_(*ν*),
  c. add every vertex pair in *N*_*G*_(*ν*) to *C* in turns to find the best triangle with maximal *f*(*C*) defined as

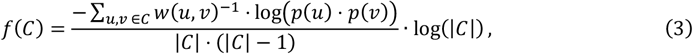
  d. where *w*(*u, ν*) denotes the Hamming distance between *u* and *ν*, and *p*(∙) is the*P*-value of a vertex; add the pair leading to the best triangle to *C*,
  e. Iteratively add neighbors of vertices in *C* one by one, until *f*(*C*) does not increase,
  f. check all vertex *ν*, in *C* excepting *ν* to see if the exclusion of *ν*, from *C* can increase *f*(∙); if yes, remove *ν*,.
4. Output all clusters ranked in decreasing order of *f*(∙).

### Motif discovery via path extension

CEMIG predicts motifs in the following three steps, digraph reconstruction, path extension, and motif refinement (**Fig. 1d**).

#### Digraph reconstruction

CEMIG reconstructs the digraph, *G*_*C*_ = (*V*_*C*_, *E*_*C*_), by merging vertices in *G*_*DB*_ based on the clusters obtained in the previous section. The vertices in the same cluster are merged into a new vertex in *G*_*C*_ named *<u>cluster vertex</u>*, and the vertices that correspond to non-significant *k*-mers are removed from *G*_*C*_. Weights of edges in *E*_*C*_are the summation of the edge weights of *G*_*DB*_

#### Path extension

CEMIG relies on a greedy algorithm to extend paths in *G*_*C*_, as follows.

1. Select an uncovered cluster with the maximum *f*(∙) as the starting vertex and initialize a path containing only this vertex,
2. For each direction (upstream or downstream), choose the edge with the maximum weight linking uncovered vertices to the path,
3. Repeat Step 2) until either of the two termination conditions occur:
  a. The path reaches length (18 − *k*),
  b. The path has been continuously added with three *k*-mer vertices in the same direction,
4. The longest sub-path composed of only the starting cluster and other cluster vertices are considered covered,
5. Output the current path and return to Step 1); if no uncovered cluster exists, output all identified paths as the seeds to refine motifs.

#### Motif refinement

CEMIG builds two sets of occurrences corresponding to the sub-path composed of the starting cluster and upstream vertices (*O*_1_), and the sub-path containing the starting cluster and downstream vertices (*O*_2_), respectively. CEMIG refines motifs as well as their lengths by building different occurrence sets based on overlaps between *O*_1_ and *O*_2_, as follows:

1. If 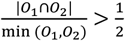, construct occurrence set *O*_1_∩ *O*_2_ of the motif corresponding to the full path.
2. If 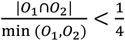, build occurrence sets, *O*_1_ *O*_2_ and *O*_1_ ∩ *O*_2_, corresponding to three motifs, respectively.
3. Otherwise, keep the occurrence sets, *O*_1_, and *O*_2_, corresponding to two motifs, respectively.

### Benchmarking motif discovery algorithms

To assess the accuracy of motif discovery algorithms on ATAC-seq data, we obtained a collection of 129 ATAC-seq experiment datasets from the Cistrome Data Browser. Our evaluation focused on comparing the performance of CEMIG against three widely used motif-finding approaches in terms of precision, specificity, ACC, and PRC metrics^2^. Additionally, we analyzed to determine the similarity between motifs identified by CEMIG and two competing methods, namely MEME-ChIP and DESSO, using ChIP-seq data. We further compared the identified motifs with those curated in the HOCOMOCO v.11 databases, employing the TOMTOM tool to calculate Q-values for similarity assessment^48,49^.

To assess the efficacy of our approach, we employed several established and widely-used motif-finding methods for comparison. These included BioProspector, a popular statistical model-based approach^13^, XXmotif, a discriminative motif prediction algorithm based on k-mer enumeration^14^, and MEME-ChIP, which integrates statistical modeling (MEME) and discriminative motif discovery (STREME)^16^. For ChIP-seq analysis, we selected MEME-ChIP as a traditional motif finding toolkit^24^, and DESSO, a state-of-the-art deep learning-based model, known for its high performance^8^. These methods were chosen to provide a comprehensive benchmarking framework and enable a thorough evaluation of our approach.

### Prediction of cell-type-specific motif

To identify differential TFBSs, we implemented a step-by-step approach to predict differentially accessible peaks between the GM12878 and K562 cell lines^25^. The process involved the following steps: (*i*) We merged all peaks obtained from the two cell lines and utilized BEDTools to count the number of reads falling into each merged peak. This was performed using the filtered alignments (BAM files) associated with the peaks. (*ii*) Optionally, we normalized the read counts to account for sequencing depth and peak length variations. This normalization step aimed to reduce potential biases and improve comparability between samples. Eq. (4) was employed for this purpose. (*iii*) To assess the significance of differences between GM12878 and K562, we employed DESeq2, a differential expression analysis tool, on the normalized read counts. (*iv*) Based on the results obtained from DESeq2, we selected peaks with a *Q*-value < 0.05 and a log2 fold-change less than −2 as GM12878-specific binding sites. Peaks with a *Q*-value < 0.05 and a log2 fold-change greater than 2 were identified as K562-specific binding sites. Shared binding sites were defined as peaks with a *Q*-value > 0.1 and an absolute log2 fold-change less than 1. Peaks that did not meet these criteria were discarded. It is important to note that the thresholds for *Q*-value and log2 fold-change can be adjusted to control the number and quality of the selected peaks, allowing for customization based on specific experimental requirements and desired stringency levels.

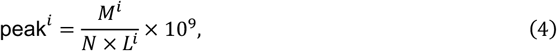

where *M, N*, and *L* denote the mapped reads of the *i*-th peak, the total mapped reads of all peaks and the length of the *i*-th peak, respectively.

## Data availability

All data utilized in this study were sourced from publicly available repositories. The ATAC-seq and ChIP-seq data employed for benchmarking purposes were retrieved using accession numbers listed in **Supplementary Table 1** and **Supplementary Table 2**, respectively. For the investigation of differential TF binding, we obtained the bed file and two replicates of bam files for the ATAC-seq dataset in GM12878 from ENCODE, specifically through the accession numbers ENCFF246XXM, ENCFF243FUG, and ENCFF832ZQV. Additionally, the bed file and two replicates of bam files for the ATAC-seq dataset in K562 were acquired from ENCODE using accession numbers ENCFF470YYO, ENCFF415FEC, and ENCFF646NWY. The RNA-seq dataset used in this case study was obtained from ENCODE, with the accession number ENCFF028CFE.

## Code availability

The C++ source code of CEMIG is freely available at https://github.com/OSU-BMBL/CEMIG.

## Acknowledgments

This work was supported by the National Key R&D Program of China (2020YFA0712400), the National Nature Science Foundation of China (NSFC, 62272270 and 11931008), and the Shandong University multidisciplinary research and innovation team of young scholars (2020QNQT017). The authors thank Dr. Yu Zhang for providing suggestions on program development.

## Author contributions

B.L. and Q.M. conceived the basic idea and designed the framework. Y.W. and Y.L. wrote the source code of CEMIG. Y.W., Y.L., and C.W. carried out benchmark experiments. Y.W. and Y.L. carried out the case study. Y.W., Y.L., and C.W. performed tool optimizations. Y.W., Y.L., Q.M., and B.L. led the figure design and manuscript writing. All authors participated in the interpretation and writing of the manuscript.

## Competing interests

The authors declare no competing interests.

## Notes

### Competing Interest Statement

The authors have declared no competing interest.

## References

1 Li, Y., Ni, P., Zhang, S., Li, G. & Su, Z. ProSampler: an ultrafast and accurate motif finder in large ChIP-seq datasets for combinatory motif discovery. Bioinformatics 35, 4632–4639, doi:10.1093/bioinformatics/btz290%J Bioinformatics (2019).

2 Zhang, S. et al. Assessing deep learning methods in cis-regulatory motif finding based on genomic sequencing data. Briefings in Bioinformatics 23, doi:10.1093/bib/bbab374 (2021).

3 Liu, B., Yang, J., Li, Y., McDermaid, A. & Ma, Q. An algorithmic perspective of de novo cis-regulatory motif finding based on ChIP-seq data. Briefings in Bioinformatics 19, 1069–1081, doi:10.1093/bib/bbx026%J Briefings in Bioinformatics (2017).

4 Ni, P. & Su, Z. Deciphering epigenomic code for cell differentiation using deep learning. BMC Genomics 20, 709, doi:10.1186/s12864-019-6072-8 (2019).

5 Ma, H., Wen, H., Xue, Z., Li, G. & Zhang, Z. RNANetMotif: Identifying sequence-structure RNA network motifs in RNA-protein binding sites. PLOS Computational Biology 18, e1010293, doi:10.1371/journal.pcbi.1010293 (2022).

6 Niu, M., Tabari, E., Ni, P. & Su, Z. Towards a map of cis-regulatory sequences in the human genome. Nucleic Acids Research 46, 5395–5409, doi:10.1093/nar/gky338%J Nucleic Acids Research (2018).

7 Machanick, P. & Bailey, T. L. J. B. MEME-ChIP: motif analysis of large DNA datasets. 27, 1696–1697 (2011).

8 Yang, J. et al. Prediction of regulatory motifs from human Chip-sequencing data using a deep learning framework. Nucleic Acids Research 47, 7809–7824, doi:10.1093/nar/gkz672%J Nucleic Acids Research (2019).

9 Li, Y. et al. A Weighted Two-stage Sequence Alignment Framework to Identify DNA Motifs from ChIP-exo Data. bioRxiv, 2023.2004.2006.535915, doi:10.1101/2023.04.06.535915 (2023).

10 Buenrostro, J. D., Wu, B., Chang, H. Y. & Greenleaf, W. J. ATAC-seq: A Method for Assaying Chromatin Accessibility Genome-Wide. 109, 21.29.21–21.29.29, doi:https://doi.org/10.1002/0471142727.mb2129s109 (2015).

11 Li, Z. et al. Identification of transcription factor binding sites using ATAC-seq. Genome Biology 20, 45, doi:10.1186/s13059-019-1642-2 (2019).

12 Bentsen, M. et al. ATAC-seq footprinting unravels kinetics of transcription factor binding during zygotic genome activation. Nature Communications 11, 4267, doi:10.1038/s41467-020-18035-1 (2020).

13 Liu, X., Brutlag, D. L. & Liu, J. S. in Biocomputing 2001 127–138 (World Scientific, 2000).

14 Hartmann, H., Guthöhrlein, E. W., Siebert, M., Luehr, S. & Söding, J. P-value-based regulatory motif discovery using positional weight matrices. Genome research 23, 181–194 (2013).

15 Bailey, T. L., Johnson, J., Grant, C. E. & Noble, W. S. J. N. a. r. The MEME suite. 43, W39–W49 (2015).

16 Bailey, T. L. STREME: accurate and versatile sequence motif discovery. Bioinformatics 37, 2834–2840, doi:10.1093/bioinformatics/btab203%J Bioinformatics (2021).

17 De Bruijn, N. G. J. P. o. t. S. o. S. o. t. K. N. A. v. W. t. A. A combinatorial problem. 49, 758–764 (1946).

18 Zheng, R. et al. Cistrome Data Browser: expanded datasets and new tools for gene regulatory analysis. Nucleic Acids Research 47, D729–D735, doi:10.1093/nar/gky1094%J Nucleic Acids Research (2018).

19 Mi, H., Muruganujan, A., Ebert, D., Huang, X. & Thomas, P. D. PANTHER version 14: more genomes, a new PANTHER GO-slim and improvements in enrichment analysis tools. Nucleic Acids Research 47, D419–D426, doi:10.1093/nar/gky1038%J Nucleic Acids Research (2018).

20 Kanehisa, M., Sato, Y., Kawashima, M., Furumichi, M. & Tanabe, M. KEGG as a reference resource for gene and protein annotation. Nucleic Acids Research 44, D457–D462, doi:10.1093/nar/gkv1070%J Nucleic Acids Research (2015).

21 Zhang, Y. et al. Model-based Analysis of ChIP-Seq (MACS). Genome Biology 9, R137, doi:10.1186/gb-2008-9-9-r137 (2008).

22 Li, G., Liu, B., Ma, Q. & Xu, Y. A new framework for identifying cis-regulatory motifs in prokaryotes. Nucleic Acids Res 39, e42, doi:10.1093/nar/gkq948 (2011).

23 Zhang, S. et al. MMGraph: a multiple motif predictor based on graph neural network and coexisting probability for ATAC-seq data. Bioinformatics 38, 4636–4638, doi:10.1093/bioinformatics/btac572 PMID - 35997564 (2022).

24 Machanick, P. & Bailey, T. L. MEME-ChIP: motif analysis of large DNA datasets. Bioinformatics 27, 1696–1697 (2011).

25 Zhang, Q. et al. Computational prediction and characterization of cell-type-specific and shared binding sites. Bioinformatics 39, doi:10.1093/bioinformatics/btac798 (2022).

26 Yang, X. & Vingron, M. Classifying human promoters by occupancy patterns identifies recurring sequence elements, combinatorial binding, and spatial interactions. BMC Biol 16, 138, doi:10.1186/s12915-018-0585-5 (2018).

27 Bradner, J. E., Hnisz, D. & Young, R. A. Transcriptional Addiction in Cancer. Cell 168, 629–643, doi:10.1016/j.cell.2016.12.013 (2017).

28 Jiang, W. G. et al. Tissue invasion and metastasis: Molecular, biological and clinical perspectives. Seminars in Cancer Biology 35, S244–S275, doi:https://doi.org/10.1016/j.semcancer.2015.03.008 (2015).

29 Schleussner, N. et al. The AP-1-BATF and -BATF3 module is essential for growth, survival and TH17/ILC3 skewing of anaplastic large cell lymphoma. Leukemia 32, 1994–2007, doi:10.1038/s41375-018-0045-9 (2018).

30 Zhang, L., Xiao, X., Arnold, P. R. & Li, X. C. Transcriptional and epigenetic regulation of immune tolerance: roles of the NF-κB family members. Cell Mol Immunol 16, 315–323, doi:10.1038/s41423-019-0202-8 (2019).

31 Chatterjee, B. et al. CD8+ T cells retain protective functions despite sustained inhibitory receptor expression during Epstein-Barr virus infection in vivo. PLoS Pathog 15, e1007748, doi:10.1371/journal.ppat.1007748 (2019).

32 Qiu, Z., Khairallah, C., Romanov, G. & Sheridan, B. S. Cutting Edge: Batf3 Expression by CD8 T Cells Critically Regulates the Development of Memory Populations. J Immunol 205, 901–906, doi:10.4049/jimmunol.2000228 (2020).

33 Hu, Y. et al. RUNX1 inhibits the antiviral immune response against influenza A virus through attenuating type I interferon signaling. Virology Journal 19, 39, doi:10.1186/s12985-022-01764-8 (2022).

34 Sekimata, M. et al. Runx1 and RORγt Cooperate to Upregulate IL-22 Expression in Th Cells through Its Distal Enhancer. The Journal of Immunology 202, 3198–3210, doi:10.4049/jimmunol.1800672%J The Journal of Immunology (2019).

35 Xu, J., Du, L. & Wen, Z. Myelopoiesis during Zebrafish Early Development. Journal of Genetics and Genomics 39, 435–442, doi:https://doi.org/10.1016/j.jgg.2012.06.005 (2012).

36 Pahl, M. C. et al. Transcriptional (ChIP-Chip) Analysis of ELF1, ETS2, RUNX1 and STAT5 in Human Abdominal Aortic Aneurysm. 16, 11229–11258 (2015).

37 Pahl, M. C. et al. Transcriptional (ChIP-Chip) Analysis of ELF1, ETS2, RUNX1 and STAT5 in Human Abdominal Aortic Aneurysm. Int J Mol Sci 16, 11229–11258, doi:10.3390/ijms160511229 (2015).

38 Teppo, S. et al. Genome-wide repression of eRNA and target gene loci by the ETV6-RUNX1 fusion in acute leukemia. 26, 1468–1477 (2016).

39 Palmi, C. et al. Cytoskeletal Regulatory Gene Expression and Migratory Properties of B-cell Progenitors Are Affected by the ETV6–RUNX1 RearrangementETV6–RUNX1 Inhibits CXCL12-Driven Cell Migration. 12, 1796–1806 (2014).

40 Hui, H. X. et al. ZNF418 overexpression protects against gastric carcinoma and prompts a good prognosis. Onco Targets Ther 11, 2763–2770, doi:10.2147/ott.S160802 (2018).

41 Luo, Y. et al. New developments on the Encyclopedia of DNA Elements (ENCODE) data portal. Nucleic Acids Research 48, D882–D889, doi:10.1093/nar/gkz1062%J Nucleic Acids Research (2019).

42 Wagner, S. et al. A broad role for the zinc finger protein ZNF202 in human lipid metabolism. J Biol Chem 275, 15685–15690, doi:10.1074/jbc.M910152199 (2000).

43 Hu, C. et al. Effects of miR-210-3p on the erythroid differentiation of K562 cells under hypoxia. 24, 1–12 (2021).

44 Skarp, S. et al. Exome sequencing reveals a phenotype modifying variant in ZNF528 in primary osteoporosis with a COL1A2 deletion. 35, 2381–2392 (2020).

45 Ma, X. et al. MiR-486-5p-directed MAGI1/Rap1/RASSF5 signaling pathway contributes to hydroquinone-induced inhibition of erythroid differentiation in K562 cells. Toxicology in Vitro 66, 104830, doi:https://doi.org/10.1016/j.tiv.2020.104830 (2020).

46 Jen, J. & Wang, Y.-C. Zinc finger proteins in cancer progression. Journal of Biomedical Science 23, 53, doi:10.1186/s12929-016-0269-9 (2016).

47 Ramos-Brossier, M. et al. Novel IL1RAPL1 mutations associated with intellectual disability impair synaptogenesis. Hum Mol Genet 24, 1106–1118, doi:10.1093/hmg/ddu523 (2015).

48 Kulakovskiy, I. V. et al. HOCOMOCO: expansion and enhancement of the collection of transcription factor binding sites models. Nucleic Acids Research 44, D116–D125, doi:10.1093/nar/gkv1249%J Nucleic Acids Research (2015).

49 Gupta, S., Stamatoyannopoulos, J. A., Bailey, T. L. & Noble, W. S. Quantifying similarity between motifs. Genome Biology 8, R24, doi:10.1186/gb-2007-8-2-r24 (2007).

