## Supplementary Information for "CEMIG: Prediction of the *cis*-regulatory motif using the De Bruijn graph from ATAC-seq"

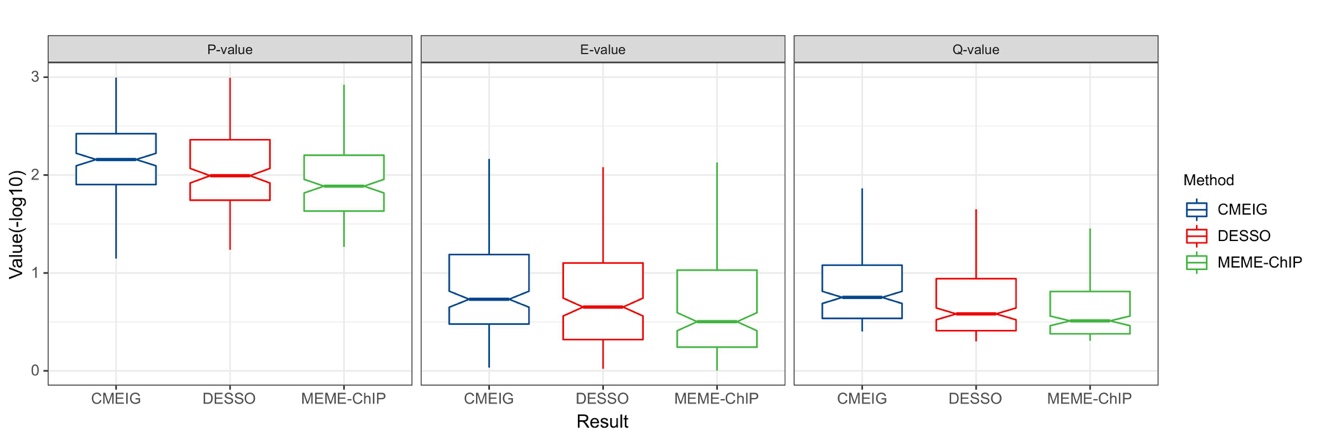

**Supplementary Fig. 1.** Benchmark motif prediction results of the 27 ChIP-seq datasets in terms of $P$-values, $E$-values, and $Q$-values of motif similarity.

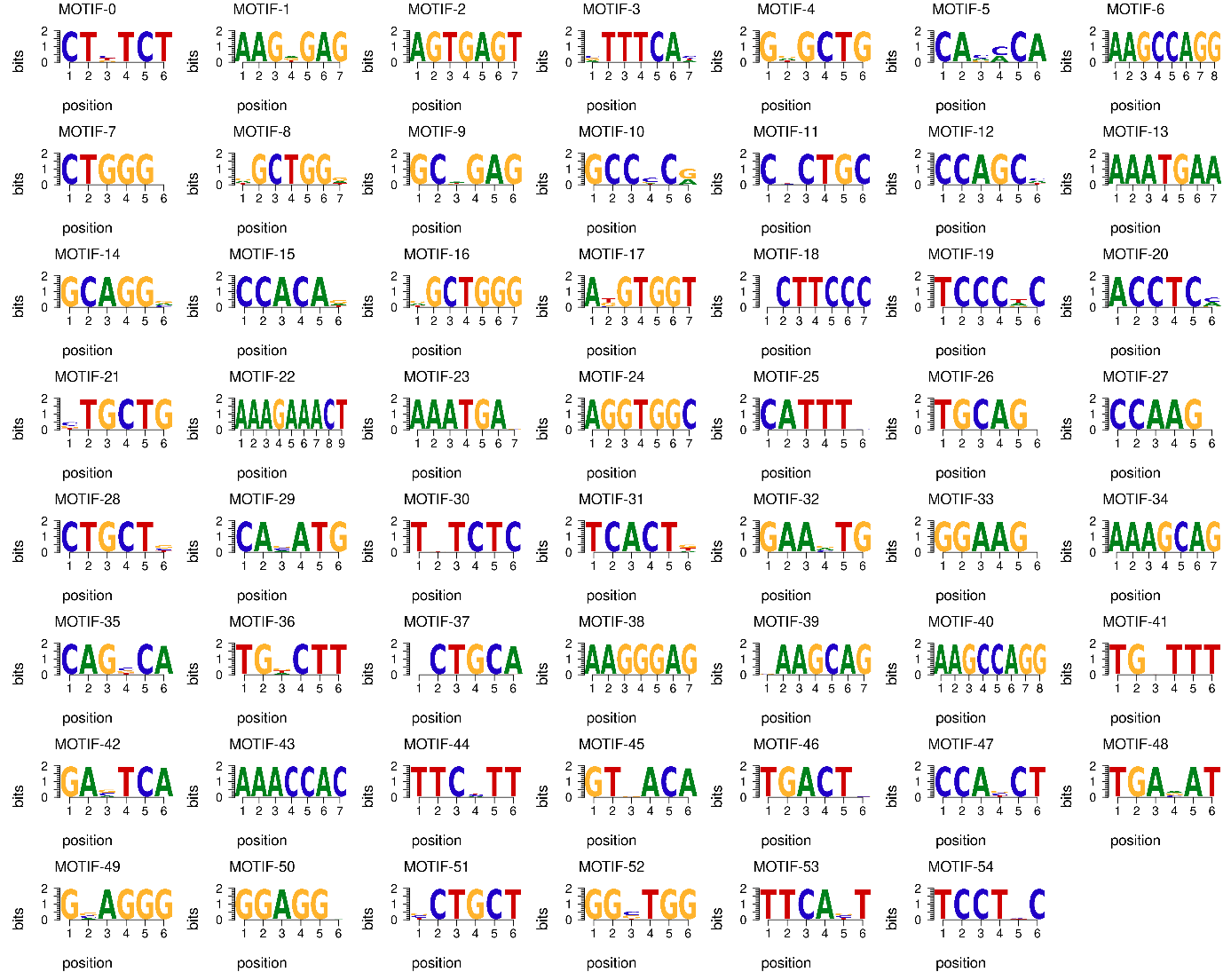

**Supplementary Fig. 2.** The sequence logos of TF motifs identified on ATAC-seq of GM12878 and K562. Motifs significantly mapped to reference motifs in HOCOMOCO or unknown ones are showcased. The correspondence relationships between discovered motifs and reference ones include: MOTIF-3 (BATF3), MOTIF-10 (AP2D), MOTIF-13 (STAT2), MOTIF-14 (ZIC2), MOTIF-15 (RUNX2), MOTIF-18 (ZN528), MOTIF-19 (NR0B1), MOTIF-21 (ZIC3), MOTIF-22 (STAT2), MOTIF-23 (NKX61), MOTIF-28 (ZIC3), MOTIF-29 (TWST1), MOTIF-32 (IRF4), MOTIF-33 (ELF5), MOTIF-36 (NR2E1), MOTIF-38 (ZNF41), MOTIF-39 (ZN418), MOTIF-41 (ZN418), MOTIF-42 (BACH2), MOTIF-43 (RUNX1), MOTIF-44 (STAT1), MOTIF-46 (BATF), MOTIF-50 (VEZF1), MOTIF-51 (ZIC3), and MOTIF-53 (IRF2).
