## Supplementary Data for "CEMIG: Prediction of the *cis*-regulatory motif using the De Bruijn graph from ATAC-seq"

**ELF3**

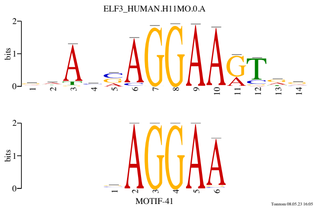


**SRR891275**

**FLI1**

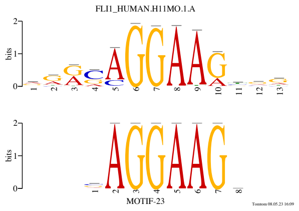


**SRR891276**

**ZNF70**

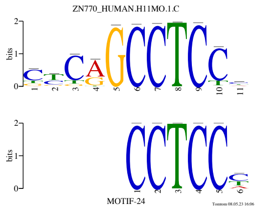


**SRR891277**

**SP1**

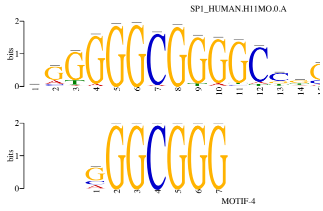


**SRR891280**

**ZN770**

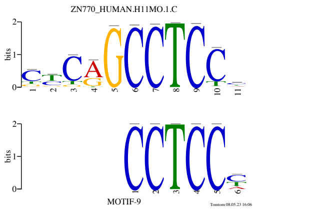


**SRR1556341**

**SP1**

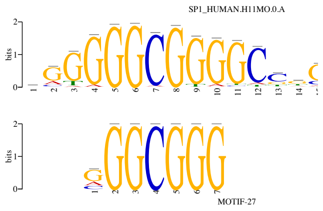


**SRR1556342**

**SP1**

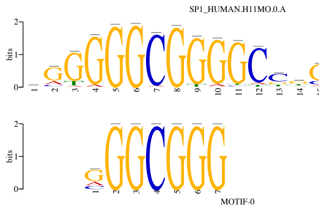


**SRR1556343**

**ZN770**

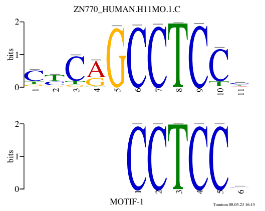


**SRR1556345**

**NFYB**

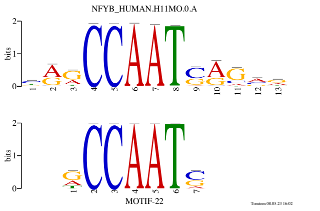


**SRR1556346**

**KLF13**

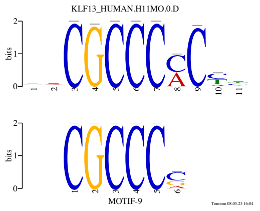


**SRR1556347**

**ZN770**

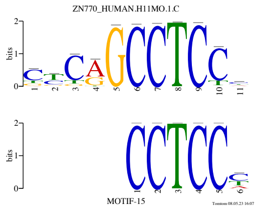


**SRR1556350**

**FLI1**

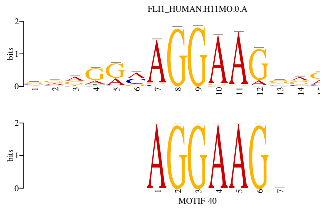


**SRR1556353**

**ZN770**

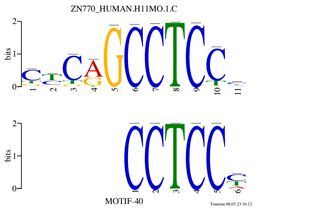


**SRR1556354**

**SP1**

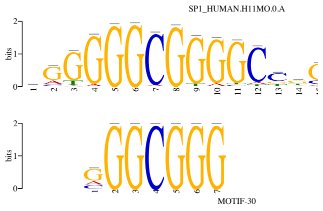


**SRR1556355**

**ZN770**

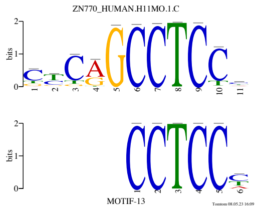


**SRR1556357**

**ZN770**

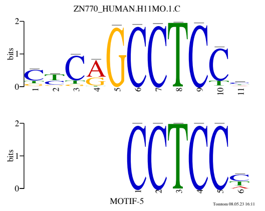


**SRR1556358**

**ELF3**

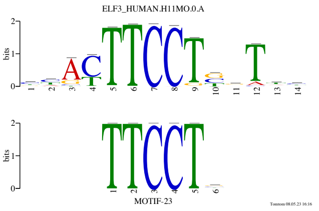


**SRR1556362**

**ZN770**

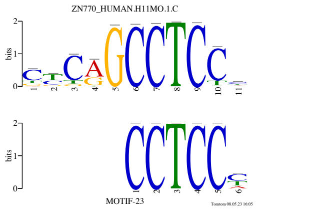


**SRR1556363**

**ZN770**

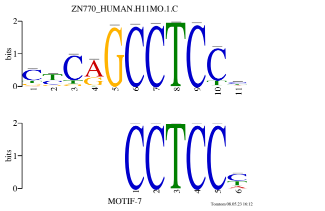


**SRR1556365**

**SP2**

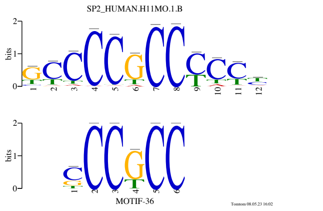


**SRR1556366**

**FLI1**

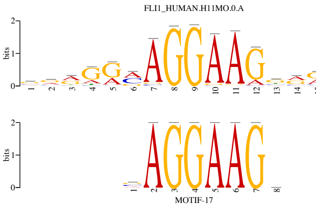


**SRR1556367**

**ZN586**

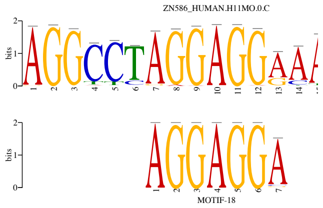


**SRR1556368**

**ZN770**

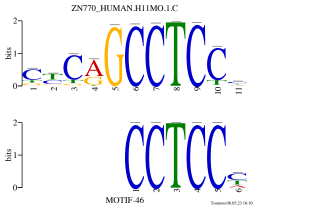


**SRR1556370**

**FLI1**

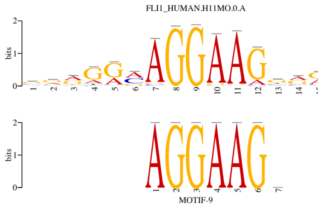


**SRR1556371**

**SP1**

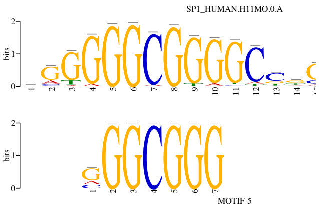


**SRR1556372**

**ZN770**

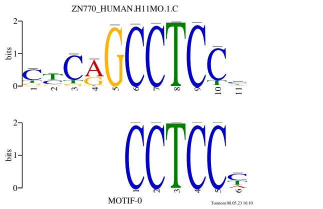


**SRR1556374**

**SP1**

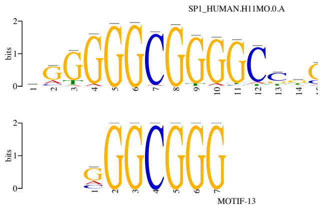


**SRR1556376**

**NFYA**

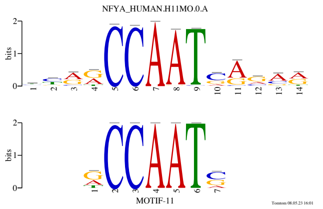


**SRR1556377**

**CR3LI**

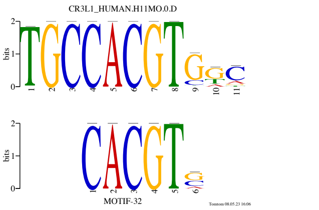


**SRR1947691**

**FLI1**

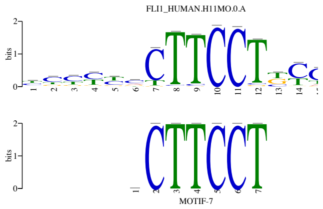


**SRR1947692**

**ZN770**

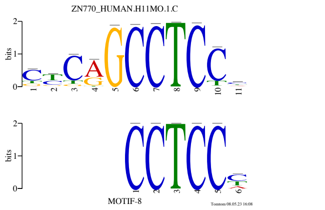


**SRR1947693**

**CEBPZ**

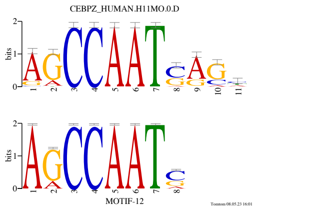


**SRR2059127**

**NFYA**

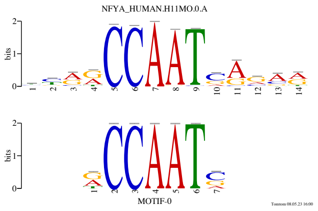


**SRR2059128**

**CEBPZ**

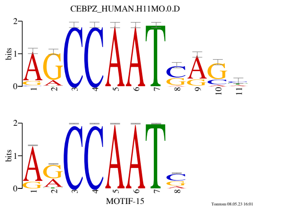


**SRR2059129**

**CEBPZ**

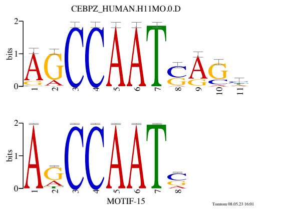


**SRR2059131**

**NFYB**

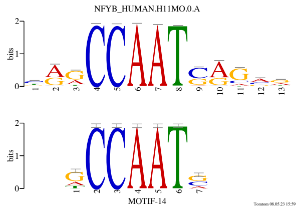


**SRR2059151**

**CEBPZ**

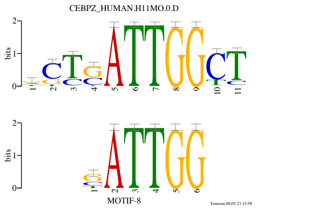


**SRR2059152**

**NFYB**

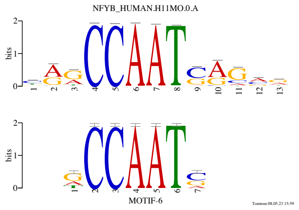


**SRR2059153**

**ATF2**

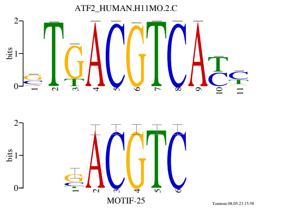


**SRR2059154**

**MLX**

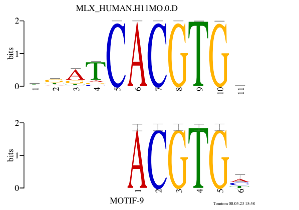


**SRR2059156**

**CEBPZ**

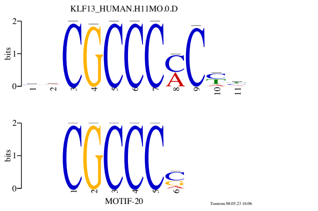


**SRR2453159**

**NFYB**

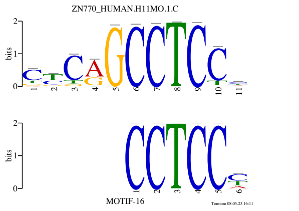


**SRR2453161**

**ATF2**

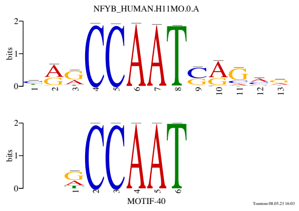


**SRR2453162**

**MLX**

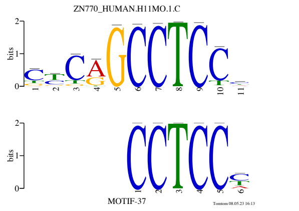


**SRR2453163**

**ZIM3**

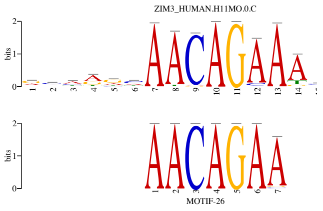


**SRR2999315**

**ZBT14**

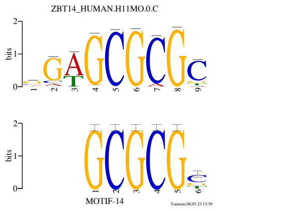


**SRR3129116**

**ELF3**

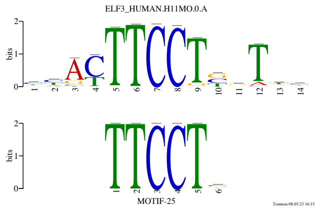


**SRR4269916**

**ZN263**

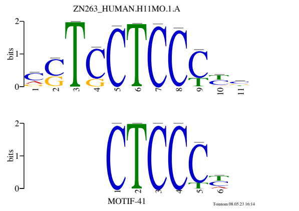


**SRR4436441**

**ZN770**

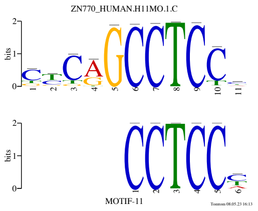


**SRR5006007**

**ZN770**

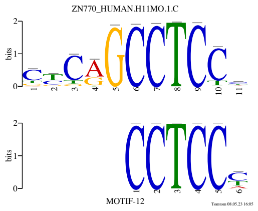


**SRR5068860**

**NFYB**

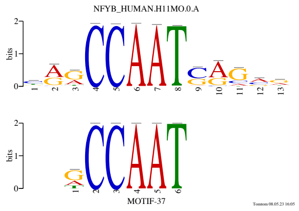


**SRR5068861**

**NFYB**

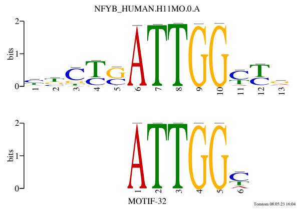


**SRR5068863**

**ZN770**

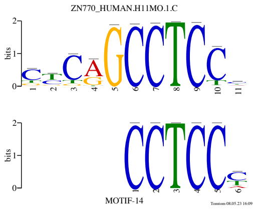


**SRR5068865**

**MAZ**

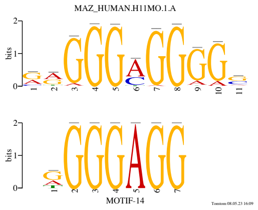


**SRR5068866**

**GLIS2**

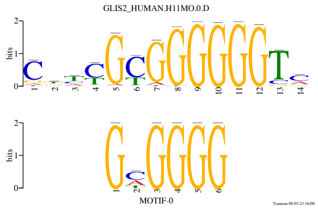


**SRR5068867**

**ZN502**

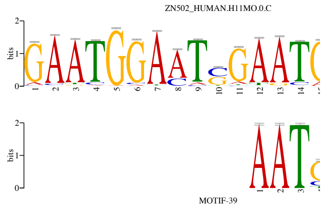


**SRR5123142**

**ZIM3**

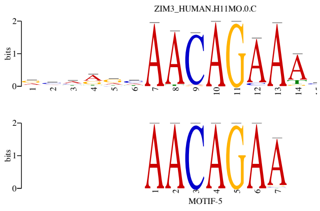


**SRR5196903**

**OLIG1**

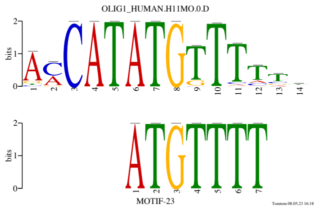


**SRR5196904**

**ATF2**

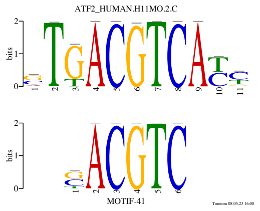


**SRR5442269**

**MAZ**

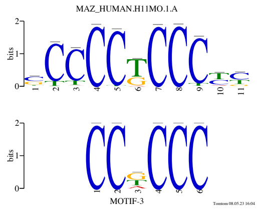


**SRR5442270**

**GLI2**

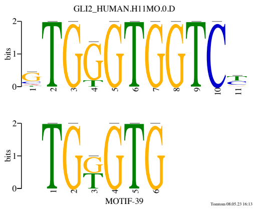


**SRR5442271**

**MAFG**

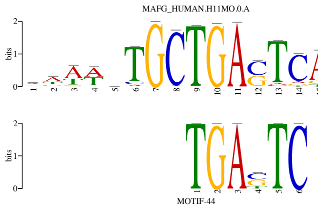


**SRR5442275**

**ZN586**

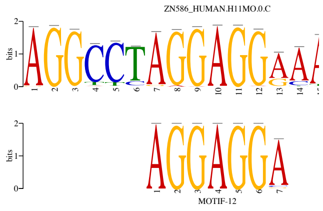


**SRR5442276**

**NFYB**

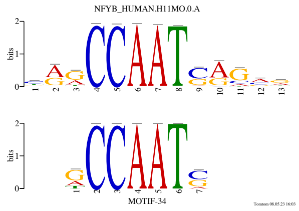


**SRR5626533**

HXB13

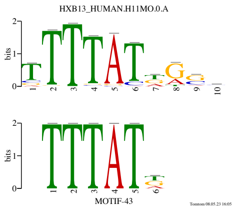


SRR5626534

NFYB

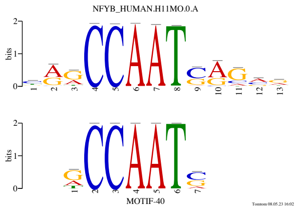


SRR5785357

MYF6

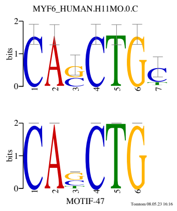


SRR5785364

ZSCA4

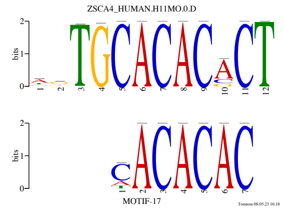


SRR5785369

FLI1

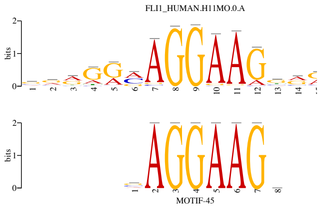


SRR5785370

NFYB

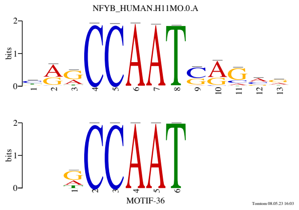


SRR5785377

ZN770

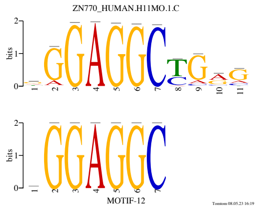


SRR5785386

FOSLI

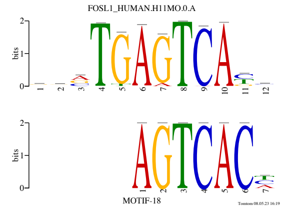


SRR5800664

MYF6

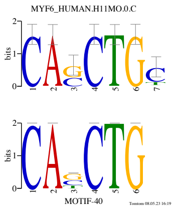


SRR5800665

ZN586

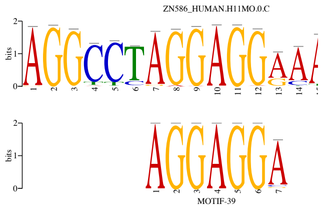


SRR5800706

SNAI2

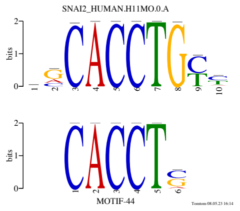


SRR5800797

ELF3

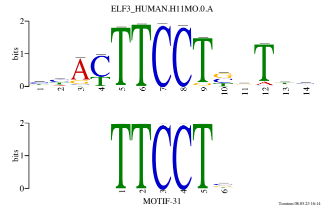


SRR5800799

MAZ

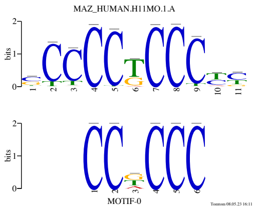


SRR5800801

GLI2

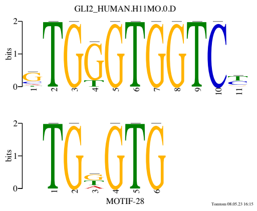


SRR5800802

ZN263

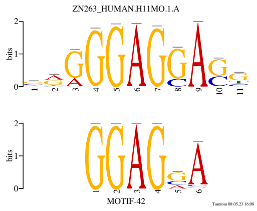


SRR6251841

ELF3

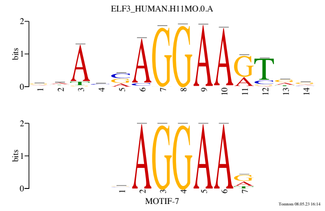


SRR6251843

**ZN770**

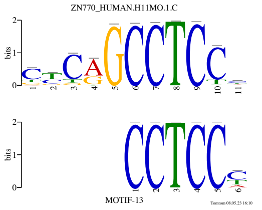


**SRR6251844**

**MAZ**

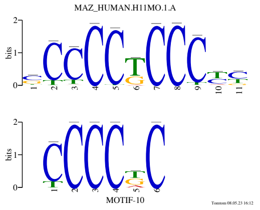


**SRR6251845**

**FLI1**

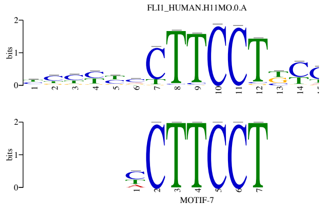


**SRR6251846**

**ELF3**

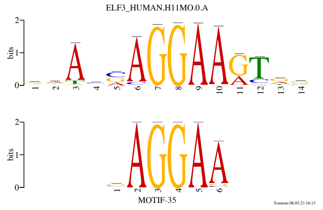


**SRR6251848**

**ZN263**

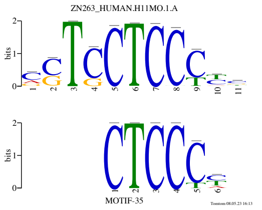


**SRR6251849**

**ZN586**

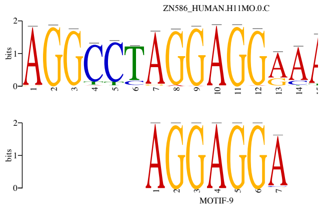


**SRR6300362**

**MAZ**

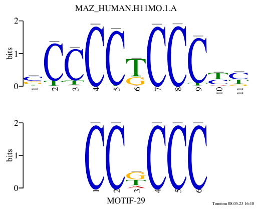


**SRR6300363**

**ETV1**

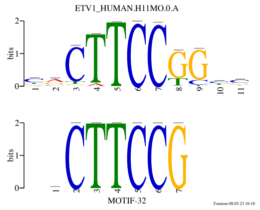


**SRR6730131**

**ELF3**

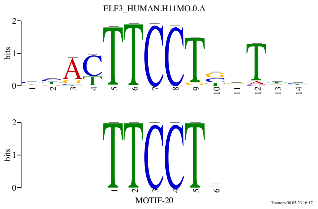


**SRR6730132**

**ZN770**

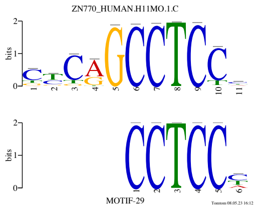


**SRR6766910**

**MAZ**

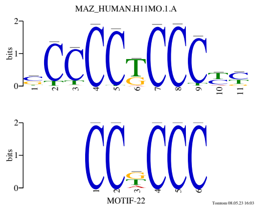


**SRR6766911**

**MAZ**

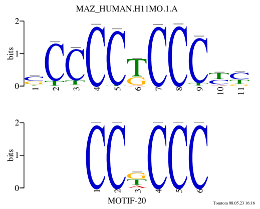


**SRR6766912**

**ZN770**

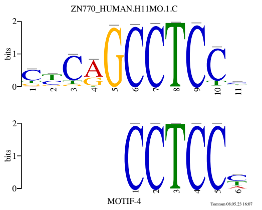


**SRR6766913**

**ZN770**

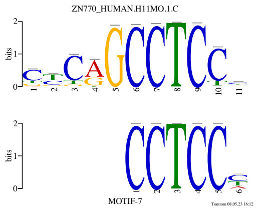


**SRR6766914**

**MAZ**

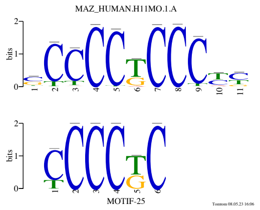


**SRR6766915**

**CPEB1**

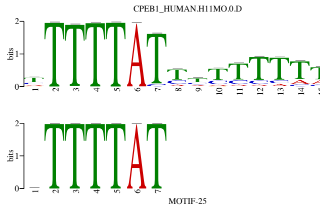


**SRR6766916**

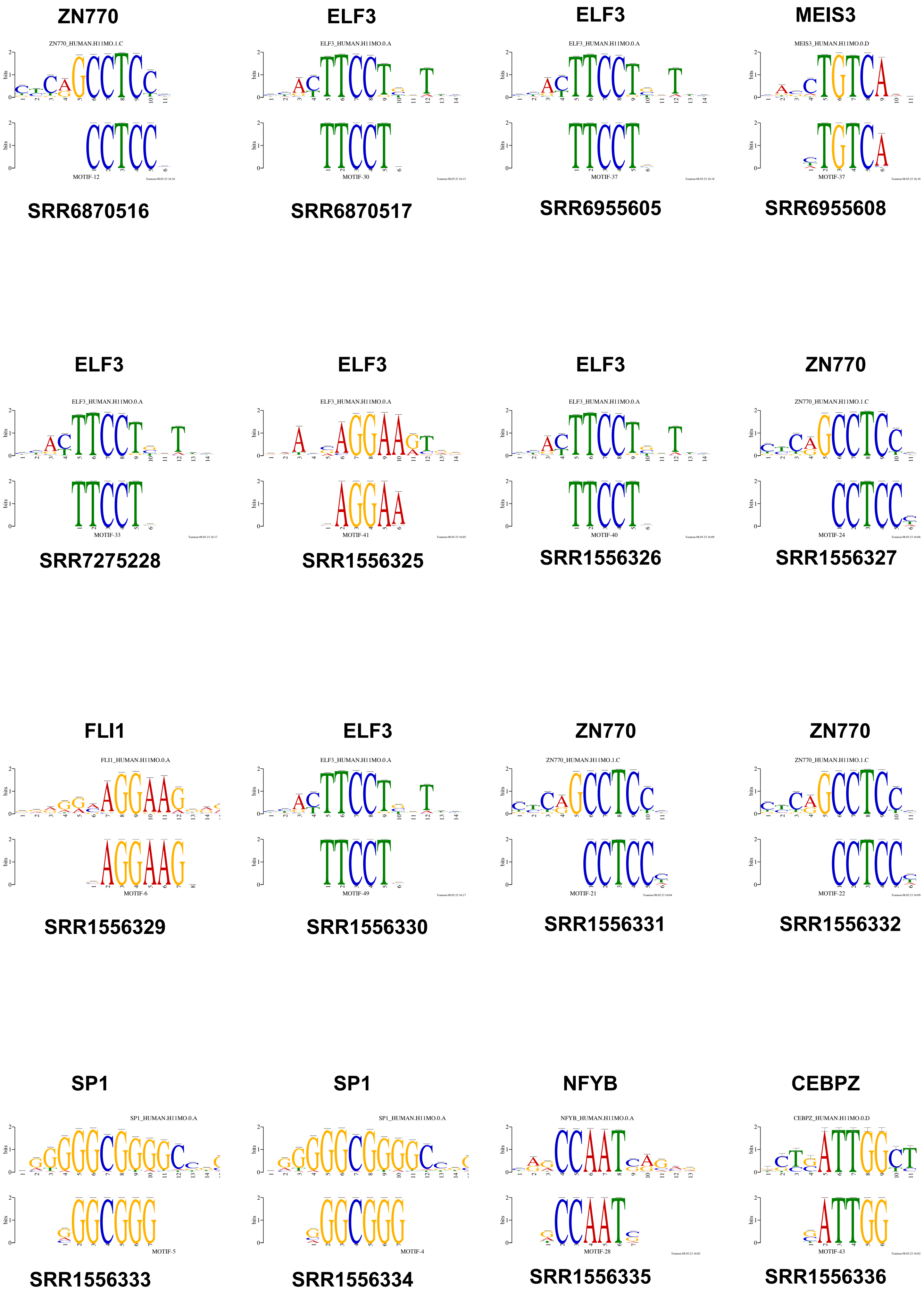

ZN770
ZN770_HUMAN.H11MO.1.C
MOTIF-12
SRR6870516
ELF3
ELF3_HUMAN.H11MO.0.A
MOTIF-30
SRR6870517
ELF3
ELF3_HUMAN.H11MO.0.A
MOTIF-37
SRR6955605
MEIS3
MEIS3_HUMAN.H11MO.0.D
MOTIF-37
SRR6955608
ELF3
ELF3_HUMAN.H11MO.0.A
MOTIF-33
SRR7275228
ELF3
ELF3_HUMAN.H11MO.0.A
MOTIF-41
SRR1556325
ELF3
ELF3_HUMAN.H11MO.0.A
MOTIF-40
SRR1556326
ZN770
ZN770_HUMAN.H11MO.1.C
MOTIF-24
SRR1556327
FLI1
FLI1_HUMAN.H11MO.0.A
MOTIF-6
SRR1556329
ELF3
ELF3_HUMAN.H11MO.0.A
MOTIF-49
SRR1556330
ZN770
ZN770_HUMAN.H11MO.1.C
MOTIF-21
SRR1556331
ZN770
ZN770_HUMAN.H11MO.1.C
MOTIF-22
SRR1556332
SP1
SP1_HUMAN.H11MO.0.A
MOTIF-5
SRR1556333
SP1
SP1_HUMAN.H11MO.0.A
MOTIF-4
SRR1556334
NFYB
NFYB_HUMAN.H11MO.0.A
MOTIF-28
SRR1556335
CEBPZ
CEBPZ_HUMAN.H11MO.0.D
MOTIF-43
SRR1556336

NFYB MAZ ZN770 ZN770

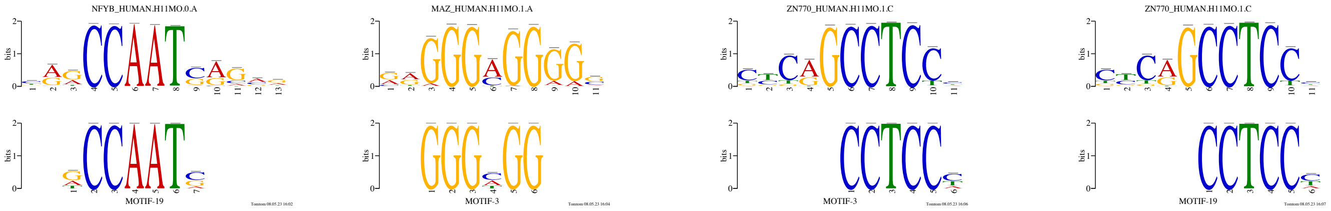


SRR1556337 SRR1556338 SRR1556339 SRR1556340
